# Integrating social-ecological dimensions of fisheries non-compliance in a stochastic framework

**DOI:** 10.64898/2026.05.05.722719

**Authors:** M. Isidora Ávila-Thieme, Kerlyns Martínez, Héctor Olivero, Mauricio Tejo, Leonardo Videla, Sergio A. Navarrete, Pablo A. Marquet, Josh Donlan, Stefan Gelcich, Rolando Rebolledo

## Abstract

Non-compliance with regulations threatens the sustainability of fisheries worldwide. Understanding the interconnected feedbacks of this complex social-ecological problem is key for sustainability but rarely integrated into fisheries management. We provide an adaptive stochastic modelling framework that integrates economic, social behavior, and ecological aspects of the Chilean kelp fishery, which plays a critical economic and ecological role in coastal social-ecological ecosystem. High levels of non-compliance is threatening sustainability, fishers’ well-being, and ecosystem health. Our model considers inherent environmental uncertainties and enables the assessment of different harvesting and compliance scenarios and the role of market-based economic incentives in reducing non-compliance. Results show that, unlike the sustainability obtained under an idealized full-compliance scenario, under dynamic compliance the social, economic, and ecological feedbacks leads to system collapse. Importantly, price premiums can promote compliance and sustainability, but the probability of collapse, albeit small, still exist. Our generalizable stochastic modeling framework evidenced that accounting for inherent uncertainty in natural resource management is key to designing interventions for sustainability.

## 1 Introduction

Achieving global sustainable fisheries presents a major challenge worldwide, especially in countries and regions of the oceans where illegal fishing, in the form of non-compliance with regulations, go unchecked and represent a significant fraction of the total biomass extracted from exploited populations [83, 1, 71, 4, 74]. These bad practices significantly contribute to the grim state of many marine resources, pushing fisheries towards paths of local or regional collapse, with the consequent degradation of human well-being and the environment [3, 72]. The illegal exploitation of species is currently classified as a wildlife crime, because it generates important economic losses, affecting peoples livelihoods, and threatening food security [81, 1]. It is therefore of paramount importance to understand the conditions that promote illegal behaviors and their consequences on sustainability, as addressing illegal fisheries is one of the goals of the 2030 Agenda for Sustainable Development. [21]. Understanding the complex interaction of social and ecological factors modulating illegal fishing can lead to rapid and long-lasting gains, including ecological, economic, and market benefits [15].

Fisheries operate as tightly coupled social-ecological systems in which human decisions and ecological processes are interconnected. Fisheries management can be decomposed into biological, environmental, social, economic, and political subsystems [38, 60]. Feedbacks between subsystems, which are key to understanding their complexity, are rarely considered in traditional fisheries science approaches [44]. Integrating the social-ecological linkages into our models can illuminate the consequences of management decisions and specific interventions available to fisheries managers [43]. Here, we used the intertidal Chilean kelp, locally known as huiro negro (*Lessonia spicata/berteroana*) fishery as a case study (henceforth intertidal kelp) to develop an open system (i.e., stochastic) model that allows us to explore social-ecological trade-offs and consequences of non-compliance with fishing regulations.

Chilean kelps play an important ecological role in coastal ecosystems [80]. They provide food and habitat for many other species [42, 55, 50, 51], are competitively dominant species in benthic ecosystems [63], contribute particulate matter (debris) to other benthic and pelagic food webs [26, 73], can store and sequester more carbon, from the atmosphere and the water column, than other macroalgae species [8], and bioaccumulate heavy metals, reducing their bioavailability and toxicity in the water column, with protective effects on other species [53, 29]. As in other temperate shores of the world, Chilean kelps also play an important economic role sustaining coastal fisheries and it is estimated that around 15,000 fishers depend directly on kelp extraction [76]. Chile leads global production of marine seaweed biomass from wild populations [54], with the kelp fishery representing between 50% and 70% of the total seaweed landings, and the intertidal kelp is the main species in the landings [66]. Most of the Chilean kelp production is exported to Asian markets for alginate extraction, a natural polymer with medical and industrial applications [25, 65, 84, 75, 46, 2]. To sustain these high levels of exploitation, the government fisheries agency (SUBPESCA) has imposed a series of science-based regulations grounded in the ecological and biological knowledge of the species, which are more strongly associated with how harvesting is conducted than with how much is harvested [76] (see table S1).

Despite regulations and their foundation on biological information, high levels of non-compliance occur in kelp fisheries, as well as in other Chilean coastal fisheries [30, 35, 24, 6]. In kelp fisheries, there are signs that kelp forests are being extracted above their natural recovery rate and that the fishery is driving the populations from mature forests to forests dominated by juveniles and to overall scarcity of plants in several sectors of the coast [34, 18]. Benthic resource fisheries operate under two management regimes in Chile: territorial user rights for fisheries (TURFs) and regional management plans (MPs). In TURFs, non-transferable area-based use rights for coastal areas are granted to organized small-scale fisher communities, who self-regulate exploitation. In contrast, MPs are regions regulated by formal instruments that establish harvesting rules and where any registered fisher is allowed to harvest [12]. These contrasting management regimes influence the overall state of the kelp forests, with apparently healthier forests inside no-take marine reserves and TURFs than in MPs, likely due to non-compliance with imposed regulations [34, 18].

The existence of two management regimes is mirrored in two types of fishers, those who are part of a fishers’ union that co-manage their resources through TURFs and non-unionized fishers whose harvesting is restricted to MPs sites. Results from previous studies have shown that those who belong to a union are more cooperative in complying with regulations because they police and punish each other. Conversely, those who do not belong to a union cooperate less with each other and are more likely to fail to comply with the regulations [33]. In this context, high levels of non-compliers can spill over to compliant fishers, undermining cooperation and triggering a breakdown of collective action [41]. Fluctuations in kelp prices, driven by international demand for kelp raw material, influence fishers’ compliance decisions by generating pulses of strong and weak exploitation intensity, with high kelp prices incentivizing fishing effort and non-compliance [58, 10]. The kelp fishery also serves as an economic entry point for the unemployed, attracting new and informal users to the fishery [10, 11]. These opportunistic fishers are less familiar with fishery regulations and have a lower sense of belonging, making them highly prone to non-compliance practices. This tendency is further motivated by fishers’ perception that enforcement of regulations is weak and the risk of being caught poaching is negligible [6].

The impacts of non-compliance with fishing regulations occur simultaneously with multiple other stressors associated with climate variability that compromise the capacity of these social-ecological systems to recover from perturbations. An example of this is El Niño Southern Oscillation events (ENSO), which can be considered as a low-frequency perturbation with important impacts on the Chilean kelp abundance. Intense El Niño events have strongly reduced the populations of these kelps, even generating local extinctions from some sites in northern Chile [16]. The ability of the population to resist and recover from these disturbances has been associated with the intensification of coastal upwelling at some sites along the shore [45]. Synoptic scale variability and high-frequency processes such as internal tides affect water temperature and nutrients delivered to the shore, and thus have significant effects on macroalgal growth rates. Thus, the inherently variable environmental conditions of the coastal ocean will affect the dynamics of kelp populations and the sustainability of the fishery. Therefore, in this study, we consider high and low frequency stochastic variability as an intrinsic feature of the system.

Market-based incentives, such as certification and eco-labelling, are considered sustainable management tools that use price premiums to discourage non-compliant fishing [62, 48, 20, 52]. However, price-based interventions can generate unintended outcomes, underscoring the need to understand the conditions under which price premiums can act as effective deterrence mechanisms. While specific deterrent strategies can encourage responsible behavior and prevent overexploitation, non-compliance is a wicked problem for fisheries worldwide, and for which there is no single silver bullet solution [59, 40, 49]. Abrupt changes in resource abundance (not only kelps), shifts in local and international market prices, and culturally-rooted behaviors among fishers represent factors that can generate or amplify divergent system trajectories leading to unsustainable and undesirable scenarios in most fisheries [39, 33, 41, 16, 31, 28, 77, 47].

There is therefore an urgent need to develop modeling frameworks that integrate the complex ecological dynamics with the multifactorial nature of fishers’ decisions. Our coupled social-ecological stochastic, kelp size-structured population model provides such a framework. We used the model to examine how uncertainty and feedbacks shape fishers’ behavior, and then assess the effectiveness of price premium deterrence strategies to reduce non-compliance (Fig. 1). By integrating ecological, fisheries and social data, the model provides an integrative framework to evaluate management-relevant interventions that foster the sustainability of complex social-ecological systems.

**Figure 1:**
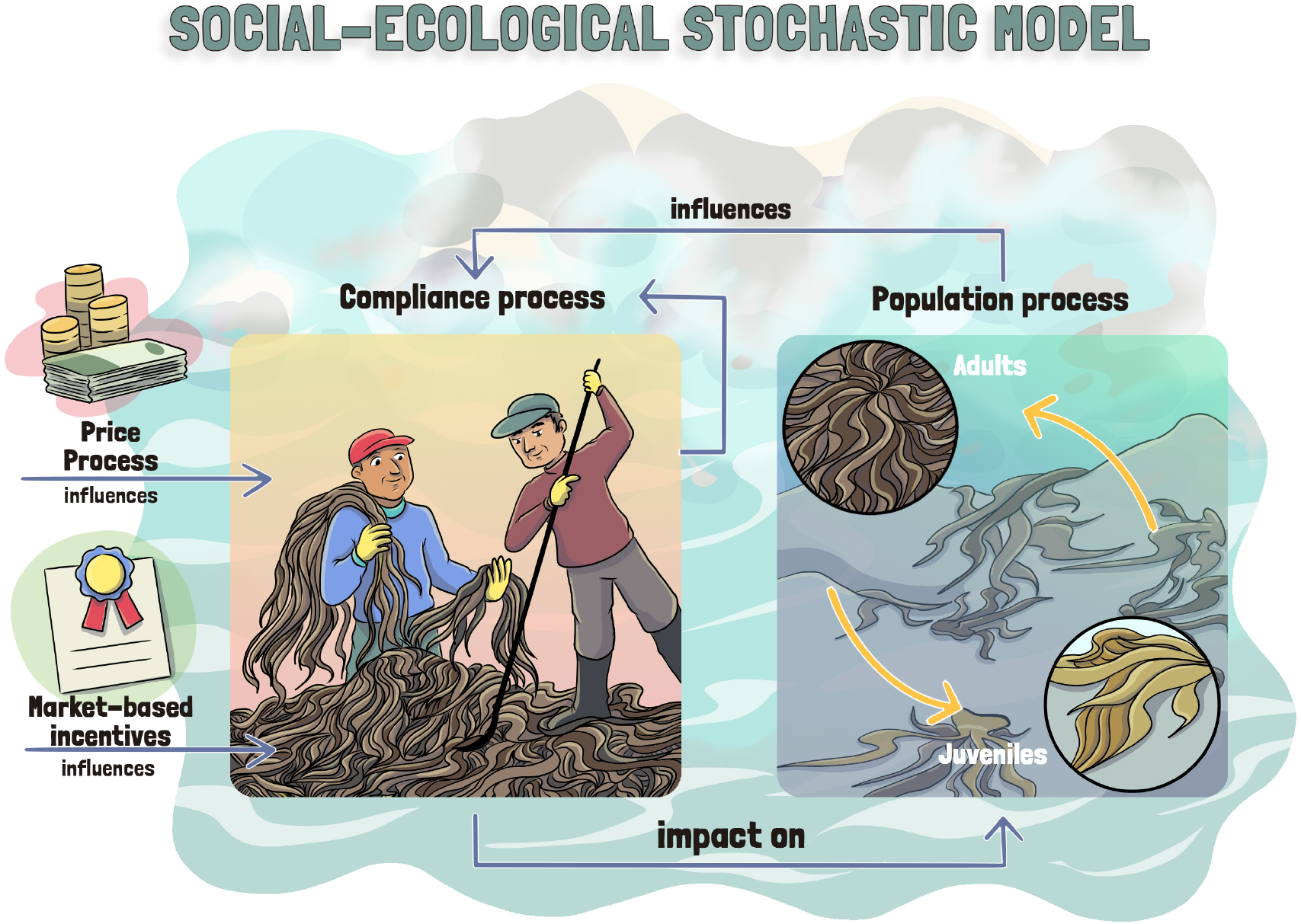
Synthesis of the component of the model and its feedbacks

## 2 Methods

### 2.1 Model description

The model we propose is composed of three components: the biological dynamics of kelp, composed of adult (*X*^(*A*)^) and juvenile (*X*^(*J*)^) stages, the fishers behavior around compliance (*E*), and the price of the kelp (*P*), which we consider as an exogenous process (Fig. 1). The model is mathematically sound, and its theoretical properties are presented in detail in [85]. In this section, we summarize the main model components, and a full description is provided in the Supplementary Material and in [85].

#### 2.1.1 The population processes

Inspired on [50], the biological dynamics of kelp is described by an age-structured model which is the target of kelp policy regulations (Table S1 in supplementary materials) and system ecology (juveniles plants provide food for several grazers, while adults plants are scarcely consumed and mainly provide habitat [42, 50]). The model is composed by the following two stochastic differential equations:

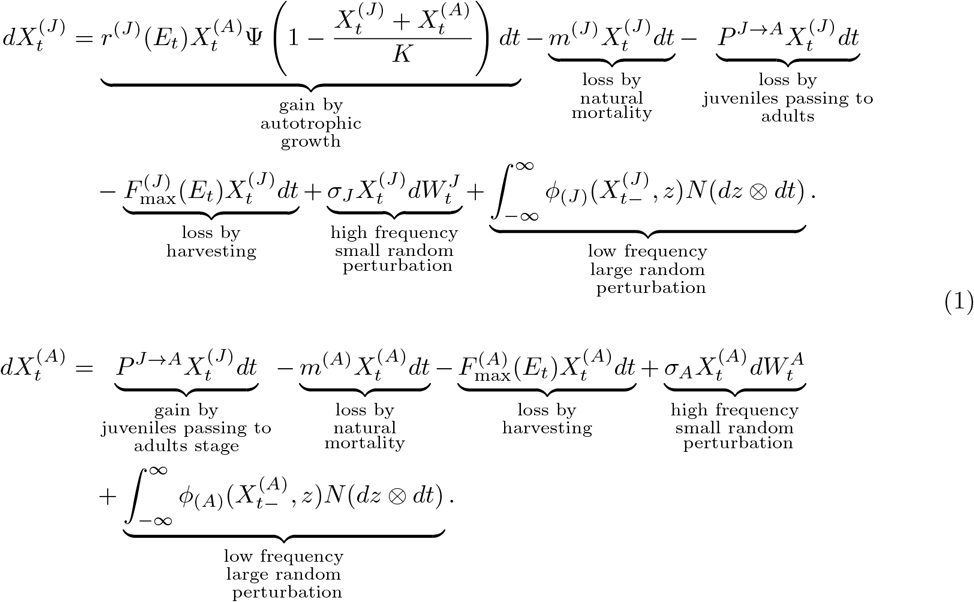

In our model, *X*^(*J*)^ and *X*^(*A*)^ represent the biomass of adults (A) and juveniles (J). The net autotrophic growth of juveniles is defined by a logistic growth function, where *r* is the intrinsic growth rate and *K* is the kelp biomass carrying capacity. As kelp is a benthic species attached to the rock surface, both stages compete for space. The parameter *m* is the fraction of biomass that is removed by non-human consumption of each stage, such as herbivory. An amount of juveniles is lost due to the probability (*P* ^*J*→*A*^) to pass from juvenile to adult, which is a positive linear function of juvenile biomass. The parameter *F*_max_ defines the fraction of biomass that is removed by harvesting. Since this model incorporates stochasticity, the logistic term must remain bounded between 0 and 1, and this is accomplished through the cut-off function Ψ, defined as follows:

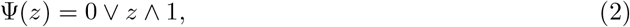

The fraction of compliant fishers (*E*) exerts direct negative effects on the adult kelp biomass 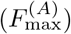, but also on the parameters associated with population recovery through juvenile growth rates (*r*^(*J*)^) and juvenile kelp biomass 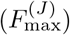 [76]. The relationship between the parameter *r*^(*J*)^ and *E* is positive; increasing compliance increases the juvenile growth rate *r*^(*J*)^ up to its maximum value defined in the model. On the contrary, the relationship between the parameter 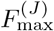 and *E* and between the parameter 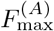 and *E* is negative, increasing compliance reduce the harvesting rate of adults and juveniles 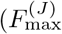 and 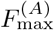.

Finally, we consider two different sources of stochasticity driving the population dynamics. The first are standard Brownian motions, *W* ^(*A*)^ and *W* ^(*J*)^, which represent frequent, small random perturbations due to, for instance, climate fluctuation and demography. The second is *N*, a Poisson random measure over [0, ∞)^2^ with intensity *ν*(*dz*) ⊗*dt* which models unfrequent but larger events, such as massive extinctions associated with ENSO events [16] (further details of the model are provided in Supplementary Material).

#### 2.1.2 The compliance process

The process *E* represents the fraction of fishers that comply with harvesting regulations. In our model, this process captures the aggregate behavior of fishers, reflecting how compliance can change over time. It is expected that *E* is in continuous feedback with the resource biomass: when the proportion of compliers decreases, extraction rates should increase, reducing resource abundance and, in turn, potentially influencing future compliance decisions. This coupling allows the model to capture the dynamic interplay between human behavior and ecosystem dynamics. The dynamic of the compliance process is given by:

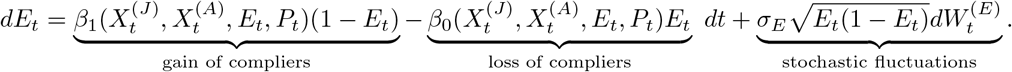

Here, the rate functions *β*_0_ and *β*_1_ define the passage from compliance to non-compliance behavior and vice versa, with *E* being negatively influenced by the proportion of fishers that change their strategies from compliance to non-compliance behavior (*β*_0_) and positively influenced by those that change it from a non-compliance to a compliance (*β*_1_). Each rate function is determined by the three decision rules associated with kelp population size (*X*), kelp price (*P*), and the fraction of fishers that have the same behavior. The decision rules are explained below, but their mathematical details are provided in the Supplementary Material 5.1.

In the first decision rule, the chance of fishers changing from compliance to non-compliance behavior increases with increasing kelp abundance, as fishers are persuaded that non-compliance will not have a major impact. On the contrary, the chance of fishers changing from non-compliance to compliance behavior increases with decreasing kelp abundance as fishers are aware that the resource is running out and that they have to behave themselves for it to recover.

In the second decision rule, the chance of fishers changing from compliance to non-compliance behavior decreases if the price is lower than a given threshold *P*_min_, at which fishers would be more willing to comply. But also, if the price is high enough, higher than *P*_max_, fishers are persuaded to not comply. We do not consider enforcement as a deterrent measure, since fishers perceive that the enforcement is weak, so they do not feel fear of being caught when non-compliant [6]. Instead, participation in a sustainability certification program may represent an alternative deterrence strategy. These programs are intended to encourage sustainable fishing practices by rewarding compliance [40, 59]. If fishers comply with harvesting regulations, they obtain an incentive associated with kelp price, which may take the form of a market price premium (s parameter in Supplementary Material). Consequently, the probability of fishers shifting from non-compliance to compliance increases when the price and the incentive exceed the compliance threshold *P*_min_.

The third decision rule is based on the fact that fishers are organized into fishers’ unions (*sindicatos*), through which they share internal rules. In cases of non-compliance with union rules, the association may exert some pressure or impose sanctions on its members. Consequently, under the influence of their union, the likelihood that a fisher changes behavior depends on the behavior of others. Here, we consider a fraction of fishers who are part of a union. Therefore, the probability of a fisher switching from compliance to non-compliance increases with an increase in the fraction of non-compliant fishers, who are members of a fishers’ union. Similarly, the chance of fishers changing from non-compliance to compliance behavior increases as the fraction of non-compliant union members rises.

The compliance process also includes a stochastic component, capturing random fluctuations in fishers’ decisions, with variability increasing as behaviors become more divided between compliance and non-compliance. This diffusion term emerges from an agent-based model where many individuals interact randomly and switch between two states (in this case, compliance and non-compliance). See the details in [85].

#### 2.1.3 The price process

The process *P* represents the kelp price, which, historically, is very variable. In some cases, the price is so low that the active extraction decreases to almost zero. Therefore, in the biological model, *P* influences *F*_max_ directly through an activation function. Here, the process *P* is modeled by a geometric Brownian motion (GBM) as used in financial practice. It has the following explicit solution:

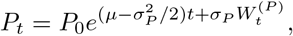

where *W* ^(*P*)^ is a standard Brownian motion accounting for random fluctuations, *µ* represents the expected return, and *σ*_*P*_ measures the volatility of the price. The initial condition *P*_0_ was established using the latest first-transaction kelp price, while the rate of return *µ* was derived from the historical first-transaction price data spanning the past 20 years [67].

### 2.2 Parameterization

All parameter values are summarized in table S2. We used a combination of empirical observations, literature sources, and survey data. The biological parameters for the process *X* were obtained mainly from previous studies [79, 50, 5]. Socioeconomic parameters for the process *X* were obtained from surveys conducted with fishers that harvest the intertidal kelp along the Antofagasta region and Atacama region of northern Chile [6]. These surveys provided estimates of compliance thresholds (*Pmin*) and initial behavioral conditions (E), whereas economic parameters such as initial prices and price dynamics were informed by historical transaction data and modelled using a Geometric Brownian Motion to account for both growth and stochastic variation. During the process of *P*, futures price thresholds were adjusted based on the average inflation of the last 30 years (1991-2022), which is 4% annually [7]. We set all model parameters on a yearly timescale. A detailed description of the parameterization procedure is provided in the supplementary materials section.

### 2.3 Simulation design

First, we simulated the kelp population dynamics (process *X*) without harvesting until the population reached a relatively steady state (10 years). At this point, we introduce harvesting simulations for another 15 years, a time at which the system reached a new steady state (Fig. 2). In harvesting simulations, we modeled two contrasting harvesting compliance scenarios: (1) full compliance and (2) dynamic compliance (Fig. 2). In the full compliance scenario, all fishers comply with harvesting regulations, meaning the process *E* is fixed at the constant level *E*_0_ = 1, and parameters related to harvesting regulations (*r*^(*J*)^, 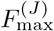, and 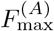) were not perturbed beyond harvesting regulations. In the dynamic compliance scenarios, the process *E* is dynamic and the parameters associated with harvesting regulations were perturbed based on the function *g*(*E, q*) detailed above (see “The population processes *X*^(*A*)^, *X*^(*J*)^”subsection).

**Figure 2:**
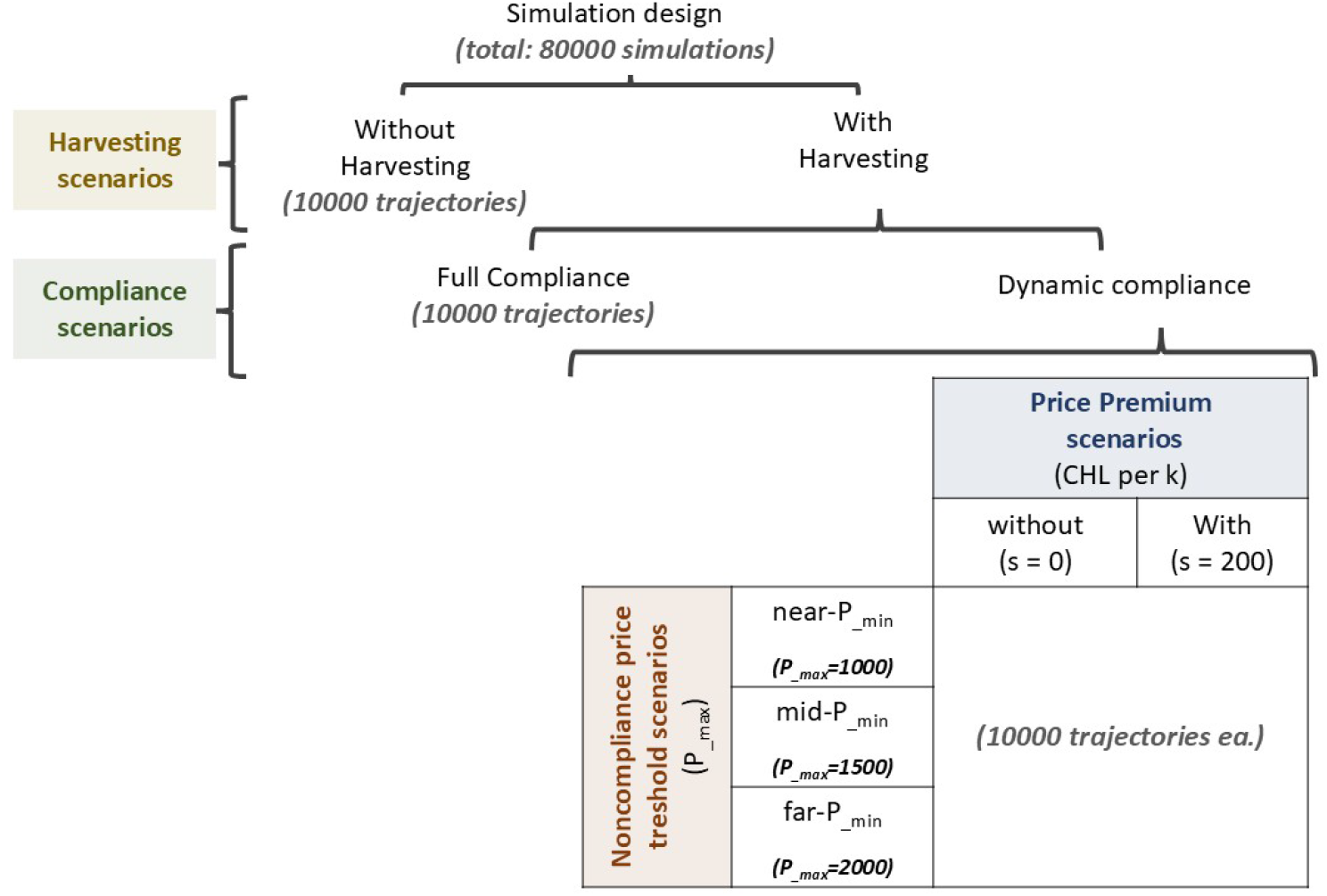
Synthesis of the design of the simulations with the number of trajectories executed in each treatment.

For dynamic compliance scenarios, we further assessed the effectiveness of a price premium in the kelp price to decrease non-compliance (i.e., without vs. with price premium) and market price threshold to non-compliance (*P*_*max*_: near-*P*_*min*_, mid-*P*_*min*_, and far-*P*_*min*_) (Fig. 2). In each treatment, only the parameter of interest was modified, while all other parameters were kept constant (*ceteris paribus*). In the price premium condition, we set the value high enough to generate a contrasting outcome, enabling us to evaluate whether this intervention could shift the fishery towards sustainable states characterized by a predominance of compliance behavior. Since *P*_*min*_ is the known minimum kelp price at which fishers are willing to comply and *P*_*max*_ the unknown price at which fishers are persuaded to not comply, smaller gaps between *P*_*min*_ and *P*_*max*_ represent greater sensitivity in switching from compliance to non-compliance behaviors.

For each price premium level, market price incentives to non-compliance and discrete shocks scenarios, we simulated 10000 trajectories. In total, we performed 260000 simulations (Fig. 2).

## 3 Results

### 3.1 Effects of Harvesting and Compliance Levels on kelp system

Under the non-harvesting scenario, the total kelp biomass was usually higher than in harvesting scenarios, stabilizing around 50000 *g/m*^2^, adding juvenile and adult biomass (see central contour in the non-harvesting scenario in Fig. 3a). On average, adults contributed approximately 60 % of the total biomass (30000 *g/m*^2^), while juveniles contributed 40 % (20000 *g/m*^2^) (Fig. 3 and Fig. S2).

**Figure 3:**
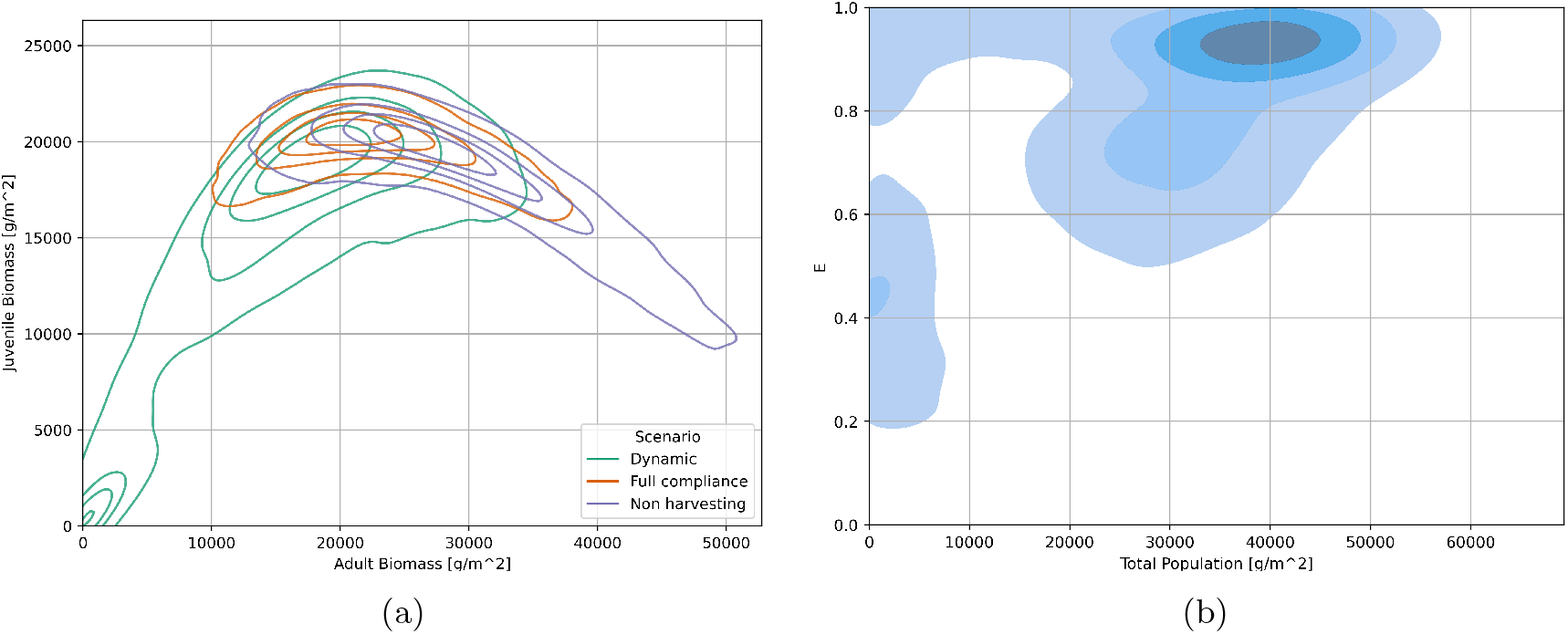
Contour plots of the probability distribution of (a) biomass of juveniles (y-axis) and adults (x-axis) evaluated at the final time step of each harvesting and compliance scenarios, and (b) the proportion of compliant fishers (*E*, y-axis) in relation to the total kelp biomass (x-axis) evaluated at the final time step of the dynamic compliance scenario. Contour lines represent different levels of the measured variable across 10000 trajectories. The innermost line corresponds to the 90th percentile, enclosing the highest values observed. Moving outward, the subsequent lines represent the 75th, 50th (median), and 25th percentiles, respectively. In (b), darker blue shades represent higher probability densities, and the innermost contours enclosing the highest values

In the full compliance harvesting scenario, the joint distribution of juvenile and adult biomass overlapped with that of the non-harvesting scenario. While adult biomass was reduced in full compliance scenarios, the population remained within sustainable states, i.e., biomass values far from zero, with both juvenile and adult biomass stabilizing at around 20000 *g/m*^2^ (Fig. 3a). This harvesting scenario resulted in a juvenile-biased population, with juveniles increasing from 40 % under non-harvesting to about 50 %, while adults decreased from 60 % to 50 % (Fig. S2).

On the contrary, the dynamic compliance scenario revealed a broader distribution of biomass values and a bimodal pattern (Fig. 3a). On the one hand, one mode corresponds to sustainable states with biomass close to the full compliance scenario. On the other hand, the second mode is located toward biomass values of juveniles and adults that approached zero, representing trajectories leading to collapsed states and highlighting the vulnerability of the system under the variable compliance behaviors. The unsustainable state with very low kelp biomass values was the most frequently observed solution, with most trajectories evolving toward total kelp population collapse (Fig. S3a).

The bimodal pattern in resulting kelp biomass also emerged with the proportion of compliant fishers (Fig. 3b). A first mode corresponded to low biomass and resulted from low to intermediate compliance fractions (*E* = 0.2 - 0.65, Fig. 3b), with most trajectories pointing toward collapse (Fig. S3a). The second mode in kelp biomass occurred under a more variable compliance pattern, where we observed trajectories with low kelp biomass and high compliance (*E* = 0.75 - 1, Fig. 3b), which represented those behaviors in which fishers switched to compliance after kelp had collapsed, i.e., waiting for recovery. These results reflect the combined influence of kelp population dynamics, social among fishers, and resource price. Across all dynamic compliance scenarios, the temporal trajectory of resource price was consistent, with most trajectories remaining between the compliance and non-compliance thresholds, while a small fraction crossed the non-compliance threshold (Fig. S3b). Thus, resource price promoted adherence to regulations when prices remained within the compliance threshold, allowing a subset of trajectories to recover partially even after kelp biomass declined. These results highlight the system’s vulnerability and emphasize the relevance of prompt compliance with sustainable harvesting practices.

### 3.2 Effect of price premium as a market-based incentive to reduce non-compliance

Without a price premium, the results mirrored those reported for the dynamic compliance scenario above. The density contours of juvenile and adult biomass and the proportion of compliers exhibited a bimodal distribution, indicating that trajectories tended to cluster around both sustainable and unsustainable states (Fig. 4a and 5a vs. Fig. 3a).

**Figure 4:**
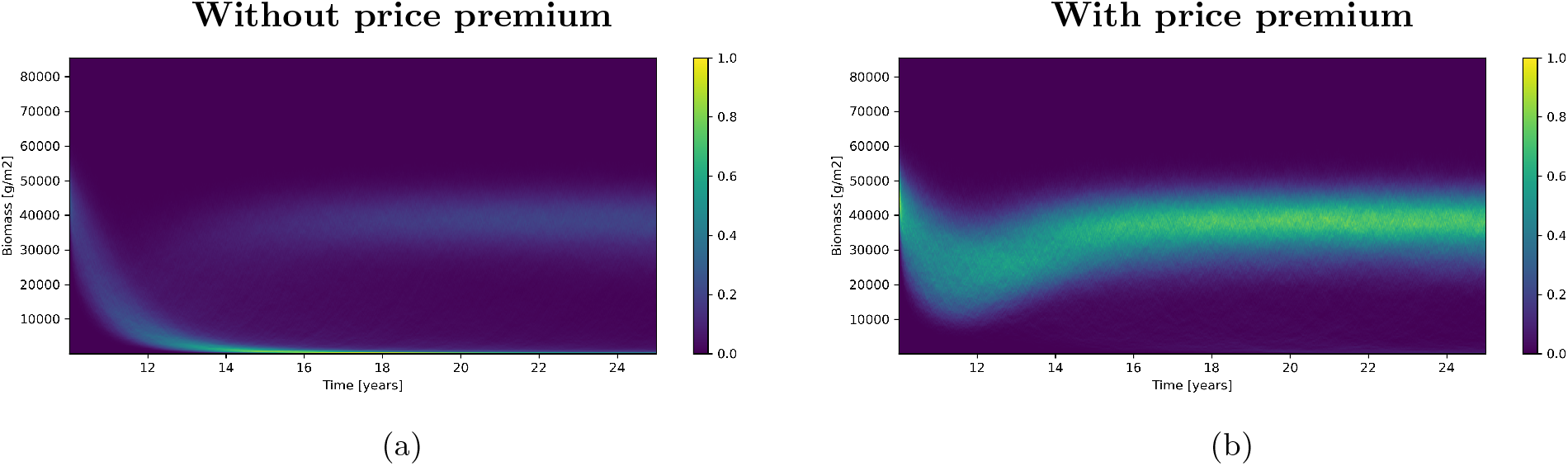
Biomass dynamics of the total kelp population over two contrasting dynamic scenarios (a) without and (b) with a price premium to the kelp price. The colour gradient represents the proportion of the 10000 trajectories occupying each state over time, with values from 0 to 1 indicating the likelihood that most trajectories are at those values at each time point

**Figure 5:**
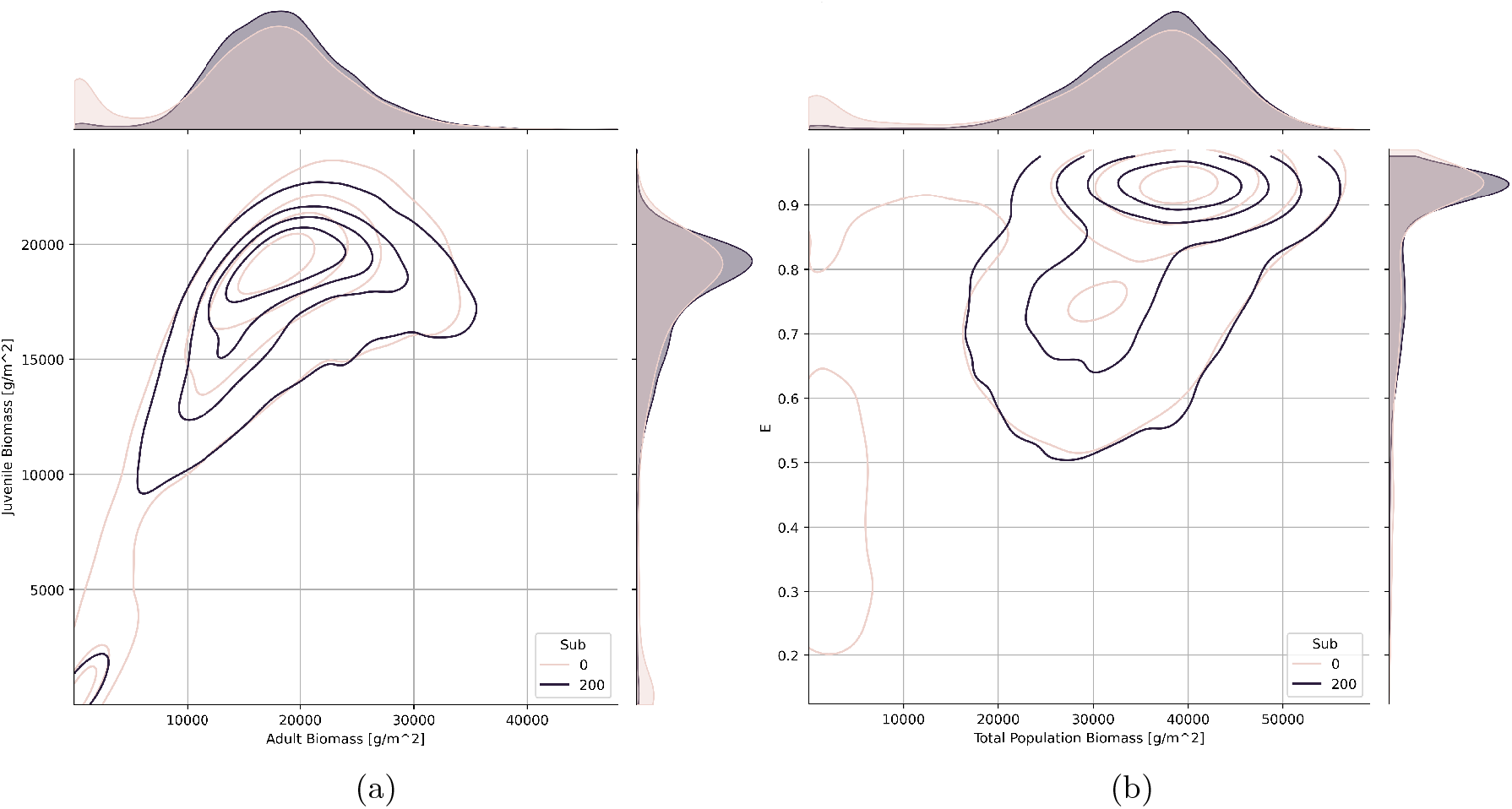
Contour plots of the probability distribution of (a) biomass of juveniles (y-axis) and adults (x-axis), and (b) the proportion of compliant fishers (*E*, y-axis) in relation to total kelp biomass (x-axis) evaluated at the final time step of each simulated trajectory. Marginal probability distributions for each axis are shown as density plots along the corresponding margins. Each panel contrasts the dynamic compliance scenario under conditions without and with a price premium (sub). Contour lines follow the same percentile interpretation described in the legend of Fig.3

When a price premium is included, the density contours moved toward sustainable zones, producing a predominantly unimodal distribution and reduced variability in both kelp biomass (Fig. 4b, 5a) and the proportion of compliant fishers (Fig. 5b). Thus, using price premium as a deterrence strategy drove the system from a regime where most of simulated trajectories resulted in kelp population collapse (Fig. 4a and 5a) toward a system in which the kelp population biomass and juveniles could persist in sustainable levels and fishers complying with harvesting regulations (Fig. 4b and 5). However, although most trajectories shifted toward sustainable states with high compliance, a small fraction of trajectories still reached unsustainable states (5a), suggesting that while the price premium substantially increases the likelihood of sustainability, a small fraction of trajectories may still lead to kelp collapse (Fig. 4, 5).

### 3.3 Effect of non-compliance price threshold (*pmax*) on dynamic compliance outcomes

The non-compliance price thresholds (*P*_*max*_) positively influenced kelp biomass and fishers’ compliance, independently of the price premium scenarios, while simultaneously reducing variability (Fig. 6). As *P*_*max*_ increased, the system tended towards higher biomass levels of juveniles and adults, and a larger proportion of compliant fishers. At the same time, increasing *P*_*max*_ reduced the variability of these ecological and social outcomes, concentrating trajectories toward the inner contour lines (Fig. 6). However, once *P*_*max*_ was above intermediate levels, further increases produced only minor changes in kelp biomass and fishers’ compliance (Fig. 6).

**Figure 6:**
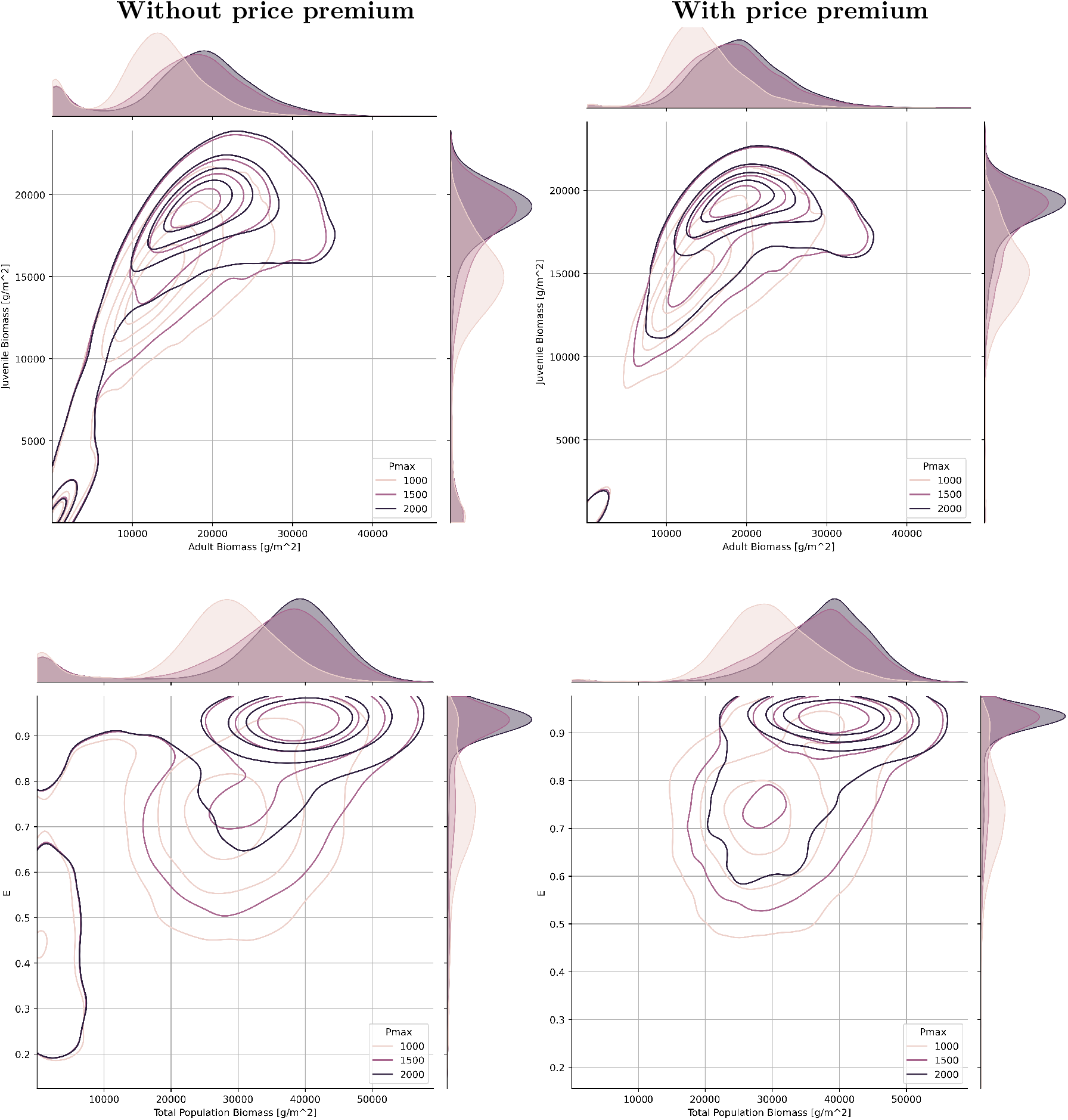
Contour plots of the probability distribution of biomass of juveniles (y-axis) and adults (x-axis, top row), and the proportion of compliant fishers (y-axis) in relation to the total kelp biomass (x-axis, bottom row) evaluated at the final time step of each simulated trajectory across three non-compliance price thresholds under dynamic compliance. Left and right column represents scenarios without and with price premiums, respectively. Marginal probability distributions for each axis are shown as density plots along the corresponding margins. Contour lines follow the same percentile interpretation described in the legend of Fig.3

## 4 Discussion

Fisheries management requires interdisciplinary approaches to consider ecological dynamics, human behavior, and socioeconomic drivers in an integrated manner, particularly when regulatory non-compliance represents a major challenge [36, 56, 1]. Here, we developed a social-ecological model that integrates these dimensions by combining empirical data with information provided by fishers to examine how feedbacks shapes compliance and sustainability in the Chilean intertidal kelp fishery, and to assess the potential impacts of deterrence interventions. Our approach provides a mechanistic understanding of the drivers of compliance and offers insights for designing interventions that promote sustainable fisheries management.

Our simulations revealed a clear gradient in the kelp forest states, ranging from mature and persistent forests under non-harvesting and full compliance scenarios to collapse under dynamic harvesting in the absence of effective deterrence strategies (Fig.3). This modeled gradient aligns with general empirical patterns reported for northern Chilean kelp beds, where forests inside notake marine reserves and TURFs appear healthier than those in management plan areas (MPs)[34]. In this sense, the empirical exploitation gradient can be conceptually interpreted as a gradient of compliance, providing an analogue for the non-harvesting, full compliance, and dynamic compliance scenarios explored in our model. While our results show that the model captures broad trends, it is not intended to predict precise outcomes, such as exact biomass values, kelp structure, and compliance levels. Rather, it provides a heuristic framework to identify underlying feedbacks and system sensitivities, drawing on empirical data and stakeholder knowledge [27, 64]. Recognizing the inherent complexity and uncertainty of social-ecological systems is key to effective management of natural resources, as it ensures that decisions are guided by an integrated understanding of ecological, social, and economic dynamics [61]. Within this context, our model contributes to the development of robust and adaptive strategies for the sustainable management of the kelp fishery.

Our study highlights that weak governance and volatile economic incentives can push the intertidal kelp system toward unsustainable states, with limited potential for recovery. This is influenced by rising kelp demand and market value over the past two decades, along with mining unemployment, fostering non-compliance, and attracting participants with weaker ties with the coastal ecosystem than traditional small-scale fishers [76, 77]. Moreover, fishers feel little to no fear of being caught as they perceive enforcement as weak or nonexistent, which promotes non-compliance [6]. The intertidal kelp fishery is therefore subject to multiple incentives for non-compliance. Strong declines in kelp populations have already been observed in MP areas, where non-compliance is more likely to occur [34, 6]. These governance and economic challenges not only undermine ecosystem resilience, but also affect the local economy, decreasing fishers’ revenues and increasing social conflicts [11]. Therefore, we highlight the need for developing effective control mechanisms to preserve the foundational role that kelps play in sustaining biodiversity, ecosystem functioning, and coastal livelihoods [23].

The deterrence strategy assessed in this study, based on economic incentives, reduced non-compliance and shifted the system away from collapse, highlighting the potential of such strategies as effective tools for fisheries management. Previous studies suggest that compliance tends to be higher when regulations target economically valuable components of the resource, such as kelp holdfasts, which are sold by weight [6]. Certification, such as being part of a sustainability program, is an important tool when dealing with non-compliance issues [59]. These programs are intended to encourage good, sustainable fishing behavior; in return, the fishery receives a price premium [40]. Examples of certification programs include the Marine Stewardship Council (MSC), which mainly certifies fish and invertebrates but has also been applied to a seaweed Korean company that obtained the world’s first certification for non-kelp cultivation [37]. However, certification programs can be expensive [17]. In Chile, the *Sello Azul* program provides a local framework to promote sustainability, ensuring the legally sourced marine resources [68]. However, this certification is primarily associated with restaurants and edible products, and therefore does not apply to kelp, which is harvested for export and industrial processing rather than direct consumption. In contrast to our results, evidence from other fisheries suggests that economic incentives do not always promote more sustainable fishing practices, as illustrated by the Norwegian haddock fishery [14]. This is likely due to higher operational costs and logistical constraints associated with maintaining catch quality and handling live fish. In a benthic kelp fishery, where harvesting is done manually and logistics are simpler, similar economic incentives may be more effective in promoting compliance and sustainable practices. Nevertheless, our results also showed that some risk of unsustainable outcomes persists even with deterrence strategies, suggesting that complementary deterrence strategies, such as strengthened enforcement, livelihood diversification, or combining formal with informal community-based mechanisms should be considered as interventions to improve sustainability. Livelihood diversification, such as nature-based tourism, can work as a deterrent mechanism by stabilizing incomes, reducing dependency on kelp harvesting, and enhancing long-term socio-ecological resilience [22, 57, 9]. Regardless of the deterrence strategy, their effectiveness also depends on other social factors such as fisher willingness, community adaptability, and local leadership [19]. Future studies should address the effectiveness of combining deterrence strategies to improve compliance.

Caveats and limitations of our study could be associated with the parametrization of our non-compliance price threshold (*P*_max_), which represents the point at which fishers’ profitability is sufficiently high that they are unwilling to comply. In our analysis, we compared three different values and observed that outcomes were sensitive to these choices. As far as we know, the price threshold that triggers non-compliance in this kelp fisheries is unknown. Future research should estimate this threshold to better inform management strategies and ensure the resilience of both the kelp population and harvesting practices. Another limitation of our study is the model used to simulate price dynamics. We adopted the Geometric Brownian Motion model for stock prices, a common approach because it is mathematically tractable (easy to analyze and simulate) and ensures that prices remain positive (values above zero). However, this model assumes that prices change at a steady proportional rate with random fluctuations, without capturing sudden shifts or market shocks. This can lead to unrealistic long-term trends, with prices growing or declining exponentially and reaching extreme values. Such behavior may not accurately reflect long-term trends in socio-economic-ecological systems, where components operate at different scales and are influenced by many external shocks. In addition, the price model is sensitive to the parameter values used. So, calibration needs to be very sharp, otherwise slight errors can lead to large errors in long-term forecasts. This is an important challenge as kelp prices are often limited at the regional and local (caleta) scale, with the most complete datasets reported as annual national averages that hide the shorter-term variability[67]. Future work should explore alternative models of price dynamics that allow for mean reversion, stochastic volatility, jumps, or memory kernels [13, 69].

## 4.1 Data and Code availability

Data and code will be made publicly available in Zenodo upon manuscript acceptance

## 4.2 Acknowledgements

We thank the artisanal fishers who participated in the surveys reported in [6], whose contributions continue to inform this study. We acknowledge to N. Segovia for his help in accessing to SECOS data center. We also acknowledge the support of Walton Family Foundation, Fondecyt 3220110, ANID PIA/BASAL, AFB240003, Millennium Science Initiative Program ICN 2019–015, and Exploration Project 13220168, who provided funding for this research. L.V. and H.O. have been partially supported by ANID through FONDECYT Iniciación 11240158 and by FONDECYT Regular 1242001, respectively. K.M. and H.O. have been partially supported by INRIA Associated Team SWAM and Programa Regional MathAmdSud AMSUD 240054.

## 5 Supplementary Material

### 5.1 Detailed model description

The model is composed of three components: the biological dynamics of kelp (*X*^(*A*)^, *X*^(*J*)^), the price of kelp (*P*), and fishers behavior around compliance (*E*). This model is mathematically well-defined, ensures well-possenes, exhibits asymptotic stability, and is suitable for numerical simulations, which means that its predictions are reliable, robust to small perturbations, and can be efficiently explored computationally [85]. Here, we provide a description of each of these processes.

#### 5.1.1 The population processes

Inspired on [50] the biological dynamics of kelp is described by an age-structured model due to its role in the kelp policy (Table S1 in supplementary materials) and system ecology (juveniles plants provide food for several grazers while adults plants mainly provides habitat [42, 50]). The model is composed by the following two stochastic differential equations:

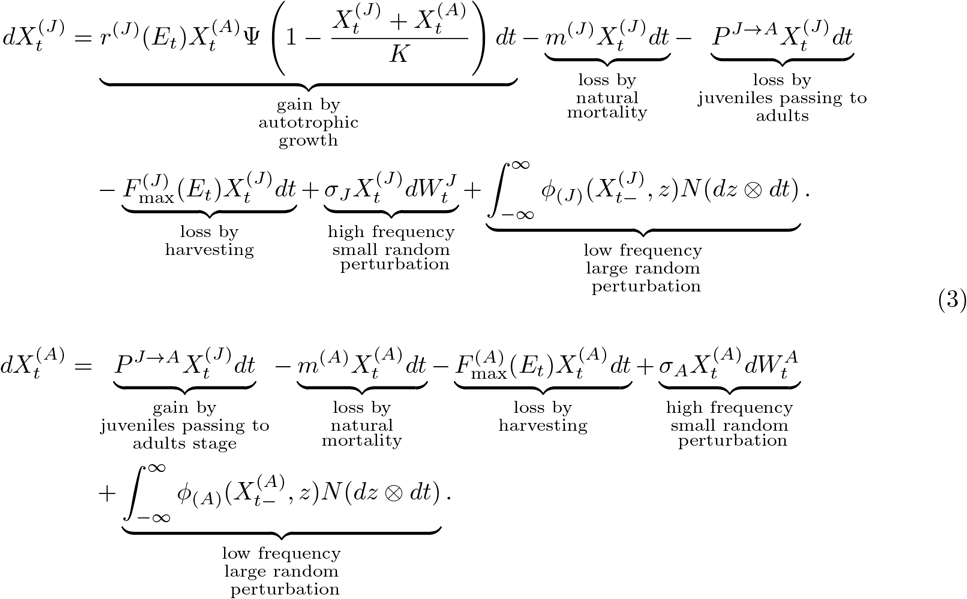

Since this model incorporates stochasticity, we need to guarantee that the logistic term remains between 0 and 1, and this is accomplished through the following cut-off function:

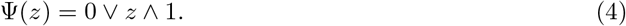

##### The deterministic terms

In the process, *X*^(*A*)^ and *X*^(*J*)^, *X*^(*A*)^ represents the biomass of adults (A), and *X*^(*J*)^ is the biomass of juveniles (J). The net autotrophic growth of juveniles is determined by a logistic growth function, where *r* is the intrinsic growth rate and *K* is the kelp biomass carrying capacity. As kelp is a benthic species, both stages compete for space. The parameter *m* is the fraction of biomass that is removed by non-human predation of each stage, such as herbivory. An amount of juvenile is lost due to the probability (*P* ^*J*→*A*^) to pass from juvenile to adult, which is a positive linear function of juvenile biomass. The parameter *F*_*max*_ defines the fraction of biomass that is removed by harvesting.

The fraction of compliant fishers (*E*) exerts direct and different effects on the parameters associated with harvesting regulations (*r*^(*J*)^, 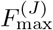, and 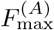) [76]. The relationship between the parameter *r*^(*J*)^ and *E* is positive, while the relationship between the parameter 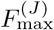 and *E* and between the parameter 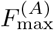 and *E* is negative. The sensitivity at which each of these parameters responds to changes in *E* is modulated by the function *g*(*E, q*) (A full description of the mechanisms describing the variation of *E* is detailed in “The compliance process”subsection).

The function *g* is defined as:

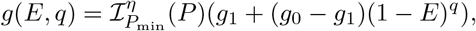

Here, *g*_0_ and *g*_1_ represent the maximum and minimum value of the function that will influence the parameters that govern harvest regulations. When there is no compliance (*E* = 0), the function reaches its maximum value 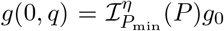. Conversely, when there is full compliance (*E* = 1), the function reaches its minimum value 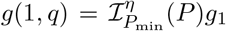. The term 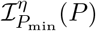 acts as an activator, it turns on this modulation only when the price *P* exceeds the threshold *P*_min_. The shape parameter *q* ∈ (0, ∞) controls whether this change from maximum to minimum happens gradually or abruptly as the fraction of compliant fishers *E* changes. Lower values of *q* produce a sharper more abrupt transition, while higher values of *q* result in a smoother, more gradual change (Fig. 7).

**Figure 7:**
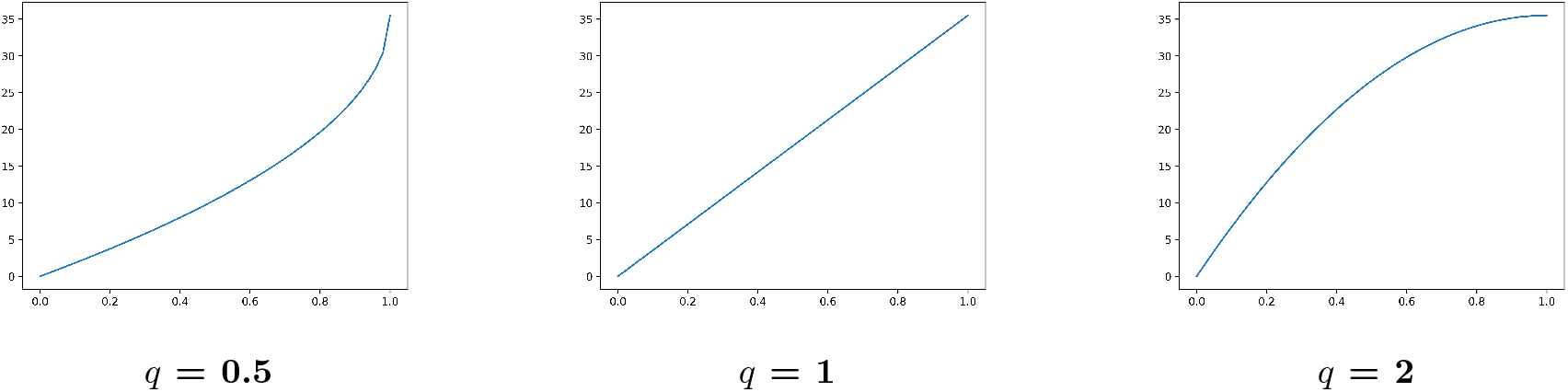
Effect of *q* values on the parameters influenced by non-compliance, in this case *rj*. X-axis represents the proportion of compliers (*E*) and y-axis the value adopted by *rj*.

##### The stochastic terms

As stated above two independent stochastic processes are driving the population dynamic: *W* ^(*A*)^ and *W* ^(*J*)^, standard Brownian motions that stand for the frequent small random perturbations, and *N* a Poisson random measure over [0, ∞)^2^ with intensity measure *ν*(*dz*) ⊗*dt* to model larger but less frequent random events, for example massive extinctions. Then, in the case of the adults, the Brownian term is

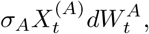

where *σ*_*A*_ is a positive parameter that modulates the size of the perturbations. Meanwhile, the Poisson term is

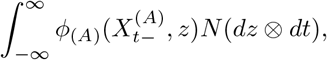

where *ϕ*_(·)_ is a response function to the jumps. In simulations we consider *ϕ*_(·)_ as a linear function.

#### 5.1.2 The compliance process

The process *E* represents the fraction of fishers that comply with harvesting regulations. In our model, this process captures the aggregate behavior of fishers, reflecting how compliance can change over time. It is expected that *E* is in continuous feedback with the resource biomass: when the proportion of compliers decrease, extraction rates should increase, reducing resource abundance and, in turn, potentially influencing future compliance decisions. This coupling allows the model to capture the dynamic interplay between human behavior and ecosystem dynamics.

The dynamic of the compliance process is given by:

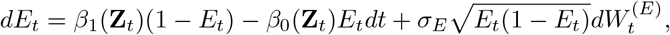

with

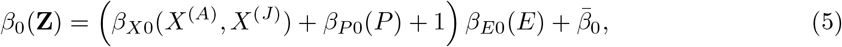

and

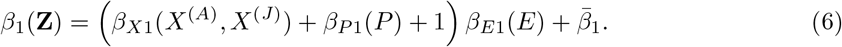

##### The deterministic terms

The rates function *β*_0_ and *β*_2_ define the passage from compliance to non-compliance behavior and vice versa, with *E* being negatively influenced by the proportion of fishers that change their strategies from compliance to non-compliance behavior (*β*_0_) and positively influenced by those that change it from a non-compliance to a compliance (*β*_2_). Each rate function is determined by the following three decision rules that are associated with the population size of the kelp (*X*), the kelp price (*P*), and the fraction of fishers that have the same behavior:

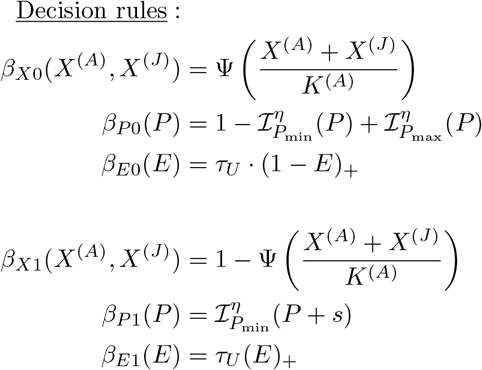

The first decision rule is associated with the population size (*X*). In this case, the chance of fishers changing from compliance to non-compliance behavior increases with increasing abundance of the kelp (*β*_*X*0_) as fishers are persuaded that non-compliance will not have a major impact. On the contrary, the chance of fishers changing from non-compliance to compliance behavior increases with decreasing abundance of the kelp (*β*_*X*1_) as fishers are aware that the resource is running out and that they have to behave themselves for it to recover.

The second decision rule is associated with the kelp price (*P*). To describe the rate functions *β*_*P* 0_ and *β*_*P* 1_ we recall the definition of Ψ in (4) to restrict the range of values between 0 and 1 and to avoid mathematical complexities. Therefore, we defined an activation function to the price rules, which is a regular version of an indicator 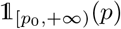:

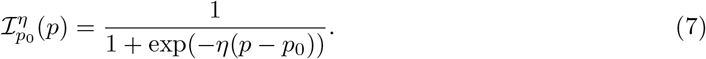

With this, the chance of fishers changing from compliance to non-compliance behavior (*β*_*P*0_) decreases if the price is lower than a given threshold *P*_min_ at which fishers would willing to comply. But also, if the price is big enough, higher than *P*_max_, fishers are persuaded to not comply. We do not consider enforcement as a deterrent measure, since fishers perceive the enforcement is weak, so they do not feel fear of being caught when non-complying [6]. Instead, participation in a sustainability certification program may represent an alternative deterrence strategy. These programs are intended to encourage sustainable fishing practices by rewarding compliance [40, 59]. If fishers comply with harvesting regulations, they obtain an incentive associated with kelp price (*s*), which may take the form of a market price premium. Consequently, the probability of fishers shifting from non-compliance to compliance (*β*_*P* 1_) increase when the price and the incentive exceed the compliance threshold 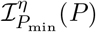. In all cases, the activation function has a sigmoid shape, whose steepness is defined by the parameter *η*.

The third decision rule is associated with the behavior of other fishers (*E*). Fishers are usually organized into fishers’ unions (*sindicatos*), through which they share internal rules. In cases of non-compliance with union rules, the association may exert some pressure or impose sanctions on its members. Consequently, under the influence of their union, the likelihood that a fisher changes behavior depends on the behavior of others (*β*_*E*0_ and *β*_*E*1_). Here, *τ*_*U*_ represents the fraction of fishers who are union members. Therefore, the probability of a fisher switching from compliance to non-compliance (*β*_*E*0_) increases with an increase of the fraction of non-compliant fishers, who are members of a fishers’ union. On the contrary, the chance of fishers changing from non-compliance to compliance behavior (*β*_*E*1_) increases as the fraction of non-compliant union members rises. Conversely, the probability of switching from non-compliance to compliance *β*_*E*0_(*E*) and *β*_*E*1_(*E*) increases as the fraction of compliant union members rises.

##### The stochastic terms

The form of the stochastic term associated to this equation,

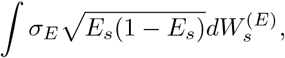

is not arbitrary; rather, it arises as a consequence of a scaling procedure applied to a sequence of coupled systems of agents that interact in a network –our fishers–, and a multi-type birth-and-death process –our kelp populations (see [85] for the rigorous mathematical details of this limiting procedure). This formulation is quite similar to the scaling-limit procedures applied to voter-like models where, in the limit, Wright-Fisher diffusions arise, with the same form of the diffusion term. Indeed, the equation for *E* can be thought of as a Wright-Fisher diffusion with bidirectional mutation (the mutation associated to the fact that fishers *mutate* their states from compliance to non-compliance and vice-versa) where the mutation rates depend also on some extra randomness (the state of juvenile and adult kelp). The explicit dependence has been explained in the previous paragraphs. It must be noticed that, with this form, we ensure that *E* remains in [0, 1], and that the variance increases as the community of fishers exhibits less consensus (alternatively, greater disagreement) on compliance/noncompliance behavior. Again, we refer the reader to [85] for the mathematical details.

### 5.2 Parameters Estimation

The supplementary Table S2 shows all the model parameters with their values and descriptions. We set all model parameters on a yearly timescale. For completeness, in the following we describe the parameterization of the three components of the model, *X, E, P*, separately.

#### For the compliance process *E*

The price threshold at which fishers are willing to comply with harvesting regulations (*P*_min_) and *τ*_*U*_ were parameterized based on empirical values obtained from surveys data that were conducted to fishers that harvest the intertidal kelp along the Antofagasta region and Atacama region of Northern Chile, between September to October 2022 [6]. These regions are the two regions that historically concentrate the largest landings at the national level [66], representing between 50% to 82% of all national seaweed landings in the last decade [66]. We surveyed fishers from Management and Exploitation Areas for Benthic Resources (MEABR) and regional management plans (MP). MEABR are Territorial User Rights Regimes (TURFs) that assign non-transferable use rights on coastal areas to organized artisanal fisher communities while MP are zones regulated by formal instruments that define the control rules for the harvesting of resources in the former de facto open access areas [35]. In total, we surveyed 196 fishers equitably from R-II (52%) and R-III (48%). In R-II, surveys were distributed in 67% from MP and 33% from MEARB. In R-III, surveys were distributed in 26% from MP and 74% from MEARB (further details details on the survey design are provided in [6]). For each harvesting regulation, we quantify the kelp price at which fishers are willing to comply with based on bidding game questionnaire [32]. The basic question was “If the price of kelp today was d, would you comply with the harvesting regulation?”. Finally, to quantify the proportion of fishers that are part of a fisher union, we asked them if they harvest from AMERBs or TURFs.

To parameterize (*P*_min_), we averaged the responses of all survey respondents (n = 196), without distinguishing between regions or management regimes. For *τ*_*U*_, we considered only the respondent who harvest from AMERBs (n= 104) and calculated the average value between the two surveyed regions. Regarding the upper price threshold at which fishers will be motivated to non-compliance (*P*_max_), the immigration into and emigration from non-compliance rates (*β*_0_ and *β*_2_), and the price premium *s* are free parameters and they were manually fitted.

#### For the price process *P*

Since we only have average prices at discrete time intervals (0, Δ, 2Δ, …, *n*Δ), we cannot observe the continuous trajectory of prices. To model their continuous evolution, we use a Geometric Brownian Motion (GBM), which captures both average return rate and stochasticity. In the Black-Scholes framework, the continuous evolution of the price of the resource *P* can be modeled as the solution of the following SDE:

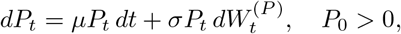

where *W* ^(*P*)^ is a standard Brownian motion accounting for random fluctuations, *µ* represents the expected return, and *σ*_*P*_ measures the volatility of the price.

The initial condition *P*_0_ was established using the latest first-transaction kelp price, while the rate of return *µ* was derived from the historical first-transaction price data spanning the past 20 years [67].

Although the GBM does not capture all the complexities of real prices, a key advantage is that its solution can be expressed explicitly as:

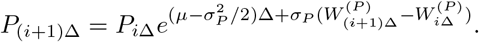

Taking expectations on both sides and using the independence of *P*_*i*Δ_ and the Brownian increments, we obtain an estimator for the average growth rate *µ*:

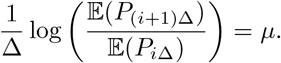

Assuming that the observed average prices are independent realizations of the GBM at each period, we have that *µ* can be estimated by

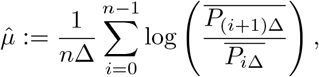

where the 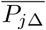’s are the observed average prices at the periods *j*Δ’s, *j* = 0, 1, …, *n*, respectively.

Under our data set, which consists of the annual average price values from the last 20 years [67], by choosing Δ = 1 year, we obtained 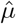. Nevertheless, the estimation for 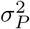 cannot be possible, since we need the information of the trajectories. Therefore, we fit 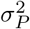 manually to represent realistic observed variations.

During the evolution of *P*, relevant parameters were adjusted to account for inflation over the last 30 years (1991-2022), which is 4% annually [7]. Parameters that were not mentioned earlier were manually calibrated since, as far as we know, no empirical estimates were available.

#### For the biomass process *X*

Most of the parameters were estimated from empirical data. For instance, the initial condition of the juvenile and adult biomasses, *X*^(*J*)^ and *X*^(*A*)^, were obtained based on empirical measures extracted from [50]. For the parameter *P* ^*J*→*A*^, we estimated the average time for a juvenile to become an adult, which was approximately one year; hence, we set *P* ^*J*→*A*^ = 1. We have assumed no ban in this fishery (*V* = 0).

For the intrinsic variances of *X*^(*J*)^ and *X*^(*A*)^ (*σ*_*A*_ and *σ*_*J*_), we first estimated the quadratic variation for the total population based on six observations that represents six years of sampling in the Chilean marine reserve “Estacion Costera de Investigaciones Marinas” (ECIM) in central south of Chile, which gives us a total estimated variability of 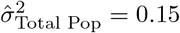. Then, we distributed this variability according to the empirical proportion of juveniles and adults of [79]:

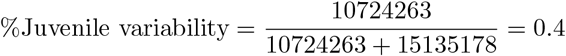

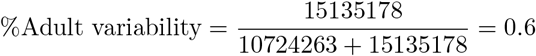

We estimate then:

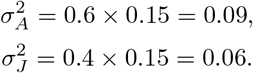

Regarding the growth rate *r*^(*J*)^, we implemented an allometric rule consistent with empirical values reported in the literature [82], obtaining the estimate *r*^(*J*)^ ≈ 35.5.

Finally, the values of *K, m*^(*J*)^ and *m*^(*A*)^ were calculated from two years of observations in kelp beds located within a marine protected area [79]. These observations measure total biomass (juveniles and adults) across seasons at a specific location in Chile, yielding a quarterly time series. Since the raw data are not directly reported, we approximated them from an image displaying the series, from which we obtained the averaged total biomass and the observed standard deviation for each observed year. Moreover, we assume that these observations were taken once the system had reached its stable state, i.e., in the absence of resource extraction or rare events, the long-term model dynamics should behave similarly. In addition, the results reported in [79] provide estimates of the biomass proportion corresponding to juveniles and adults, allowing us to translate total population data into juvenile and adult subpopulations.

Under these assumptions, we estimated the carrying capacity *K* as the maximum average biomass observed:

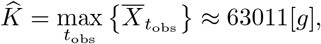

where 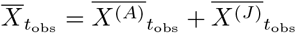 denotes the observed averaged (total) biomass at time *t*_obs_.

Finally, for the estimation of the death rates *m*^(*A*)^ and *m*^(*J*)^, we adjust the values of 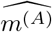 and 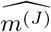 in order to find the values that best replicate the intervals reported in [79]. To do this, we simulate *M* trajectories for the kelp system without jumps and without extraction:

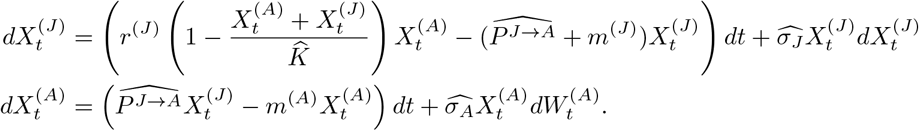

We set the initial condition as the mean biomass at equilibrium (considering the *N*_obs_ = 4 2 +1 observations):

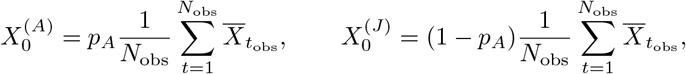

where *p*_*A*_ ≈ 0.88 represents the proportion of adult biomass observed.

Then, we compute an approximated solution for the minimization problem:

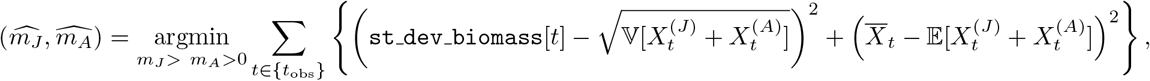

where the constraint *m*^(*J*)^ *> m*^(*A*)^ *>* 0 reflects the biological assumption that juveniles/recruits suffer high mortality due to grazing and shading, while adults tend to be more persistent [78, 50]. The minimizer is approximated numerically using standard optimization routines (for an implemented version we refer to https://github.com/holivero/Kelp). Results of the fitting can be visualize in Figure S1 below for estimates 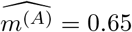 and 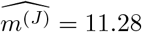.

### 5.3 Suppplementary Figures

**Figure S1:**
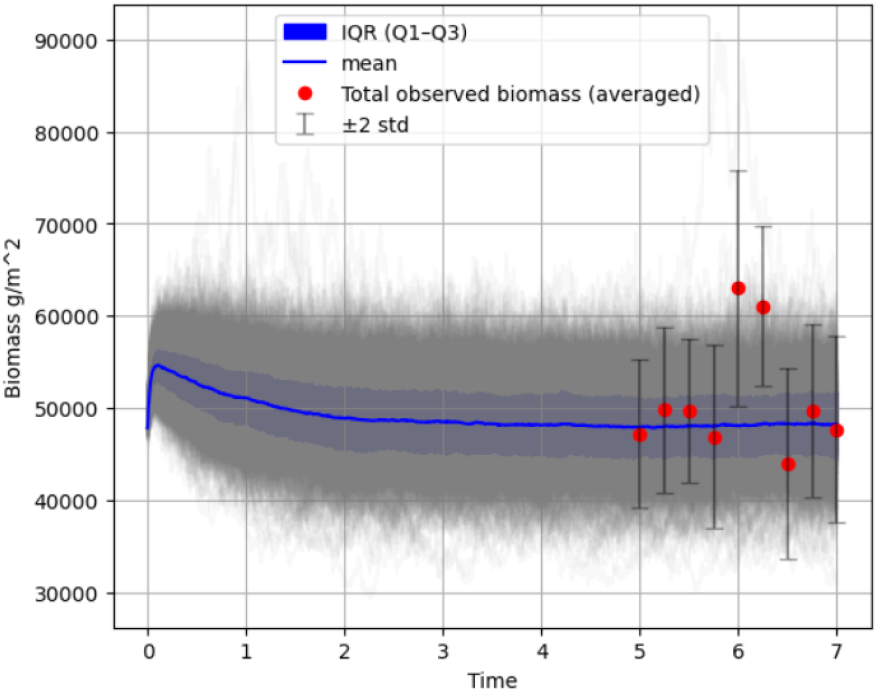
Simulated trajectories of total biomass (*g/m*^2^) under the fitted model (grey lines), with mean trajectory (blue line) and interquartile range (blue shaded area). Red dots represent averaged observed biomass per season with ±2 standard deviations (error bars).

**Figure S2:**
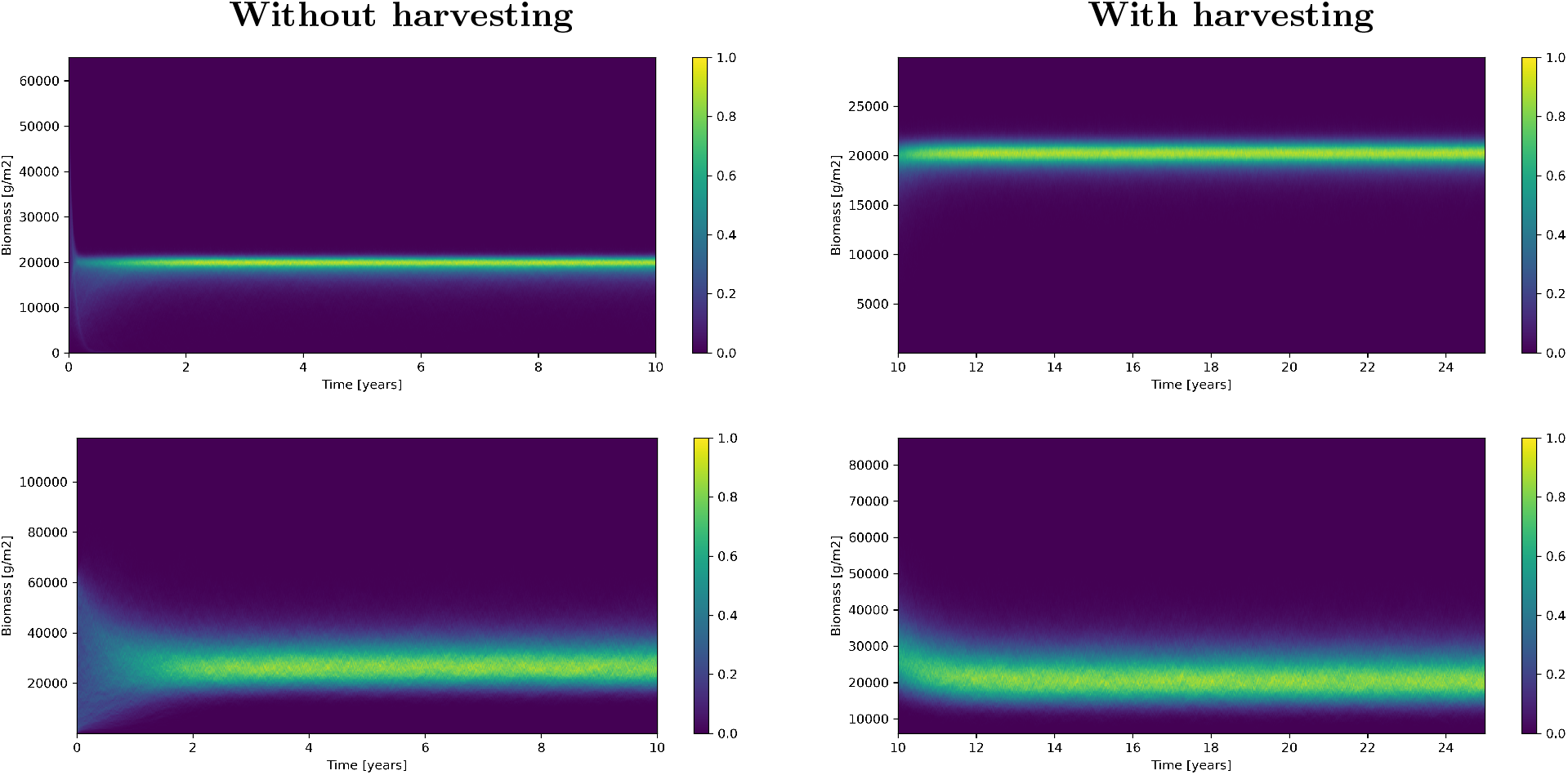
Biomass dynamic of juveniles (top row) and adults (bottom row) across time under no harvesting (left column) and harvesting (right column) scenarios. The colour gradient represents the proportion of the 10000 trajectories occupying each state over time, with values from 0 to 1 indicating the likelihood that most trajectories are at those values at each time point. Note that the y-axis is on a different scale

**Figure S3:**
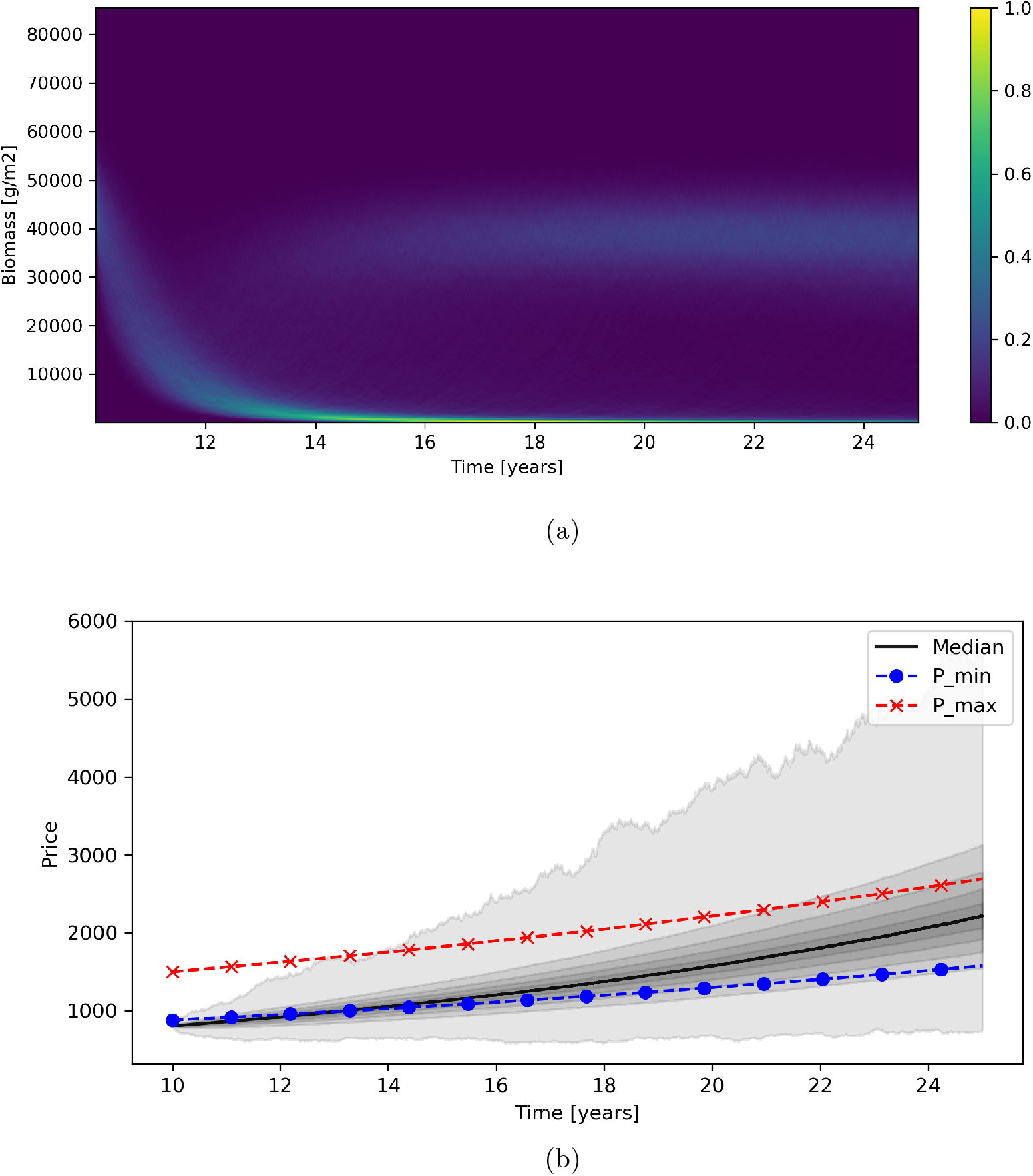
Results of the dynamic compliance scenario showing (a) Biomass dynamic of total kelp population over time and (b) kelp price (CHL per kg). In (a) trajectory colors following the same gradient as Fig. S2, ranging from blue to yellow, which indicate the likelihood that most trajectories occupy the corresponding biomass at each time point.

### 5.4 Suppplementary tables

**Table S1:** Harvesting rules of the kelp fishery and their role on the kelp population model. Parameter values and references are detailed in.

| Harvesting rule | Biological role of rule | Parameter perturbed | Compliance | non-compliance |
| --- | --- | --- | --- | --- |
| Remove only adult plants | Allows juveniles to grow and reproduce | $F_{\max}^{(J)}$ | Fishers harvest only adults | Fishers harvest adults and juveniles |
| Remove the thallus, including the holdfast to facilitate recruitment | If pruned, the plant does not regenerate. The holdfast remains in the space until died ($\sim 6$ months), During this period, recruitment is not allowed in that occupied space, decreasing the population growth rate | $r^{(J)}$ | Fishers remove entire plants | Fishers prune the plant |
| Remove one out of every three plants | To facilitate recruitment. This plant has low dispersion distance. Therefore, this distance represents the threshold to facilitate recruitment. Also, this distance allows kelp to form a forest-like ecosystem. If distances increase, even that recruitment could happen, the growth of recruits is hard due to the strong environment conditions | $r^{(J)}$ | Fishers leave sparse adults in the field | Fishers leave large barren areas |
| Do not harvest more than 25% of adult biomass | To allow population recovery after harvesting | $F_{\max}^{(A)}$ | Fishers leave $\sim 75\%$ of biomass | Fishers over-harvest the standing stock |
| Harvest only in areas with density $> 1$ plant/m<sup>2</sup> | Inspired by rule 3, avoids barren areas; low density reduces recruitment | $r^{(J)}$ | Fishers harvest in dense zones | Fishers harvest in sparse zones |

**Table S2:**
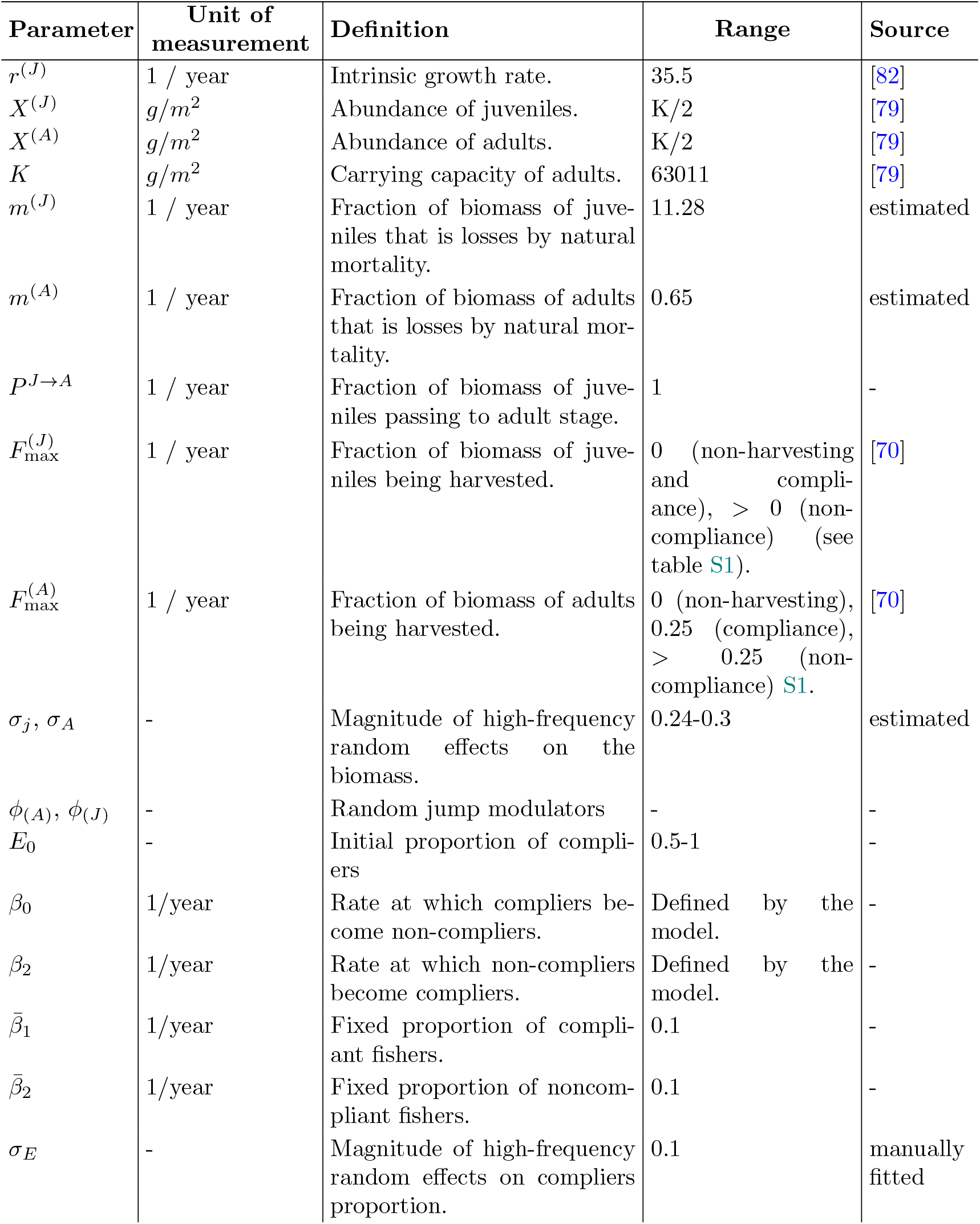
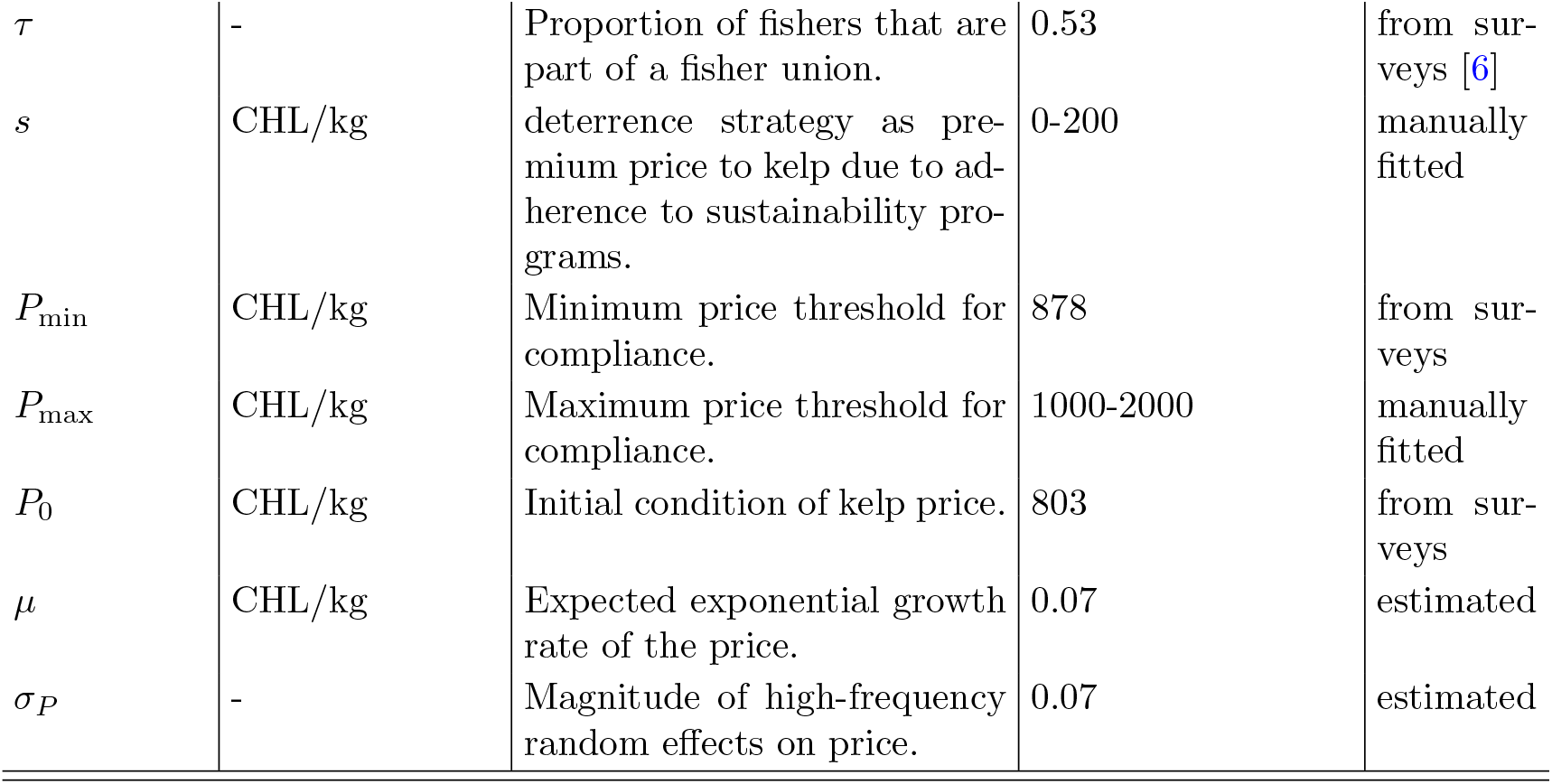
Parameter values of the kelp model.

